# Brain Age Analysis and Dementia Classification using Convolutional Neural Networks trained on Diffusion MRI: Tests in Indian and North American Cohorts

**DOI:** 10.1101/2024.02.04.578829

**Authors:** Tamoghna Chattopadhyay, Neha Ann Joshy, Saket S. Ozarkar, Ketaki Buwa, Yixue Feng, Emily Laltoo, Sophia I. Thomopoulos, Julio E. Villalon, Himanshu Joshi, Ganesan Venkatasubramanian, John P. John, Paul M. Thompson

**Affiliations:** Imaging Genetics Center, Mark and Mary Stevens Neuroimaging and Informatics Institute, Keck School of Medicine, University of Southern California, Marina del Rey, CA, United States; Multimodal Brain Image Analysis Laboratory, National Institute of Mental Health and Neuro Sciences (NIMHANS), Bengaluru, Karnataka, India; Translational Psychiatry Laboratory, National Institute of Mental Health and Neuro Sciences (NIMHANS), Bengaluru, Karnataka, India

**Keywords:** Magnetic Resonance Imaging, Diffusion Tensor Imaging, Deep Learning, Harmonization

## Abstract

Deep learning models based on convolutional neural networks (CNNs) have been used to classify Alzheimer’s disease or infer dementia severity from T1-weighted brain MRI scans. Here, we examine the value of adding diffusion-weighted MRI (dMRI) as an input to these models. Much research in this area focuses on specific datasets such as the Alzheimer’s Disease Neuroimaging Initiative (ADNI), which assesses people of North American, largely European ancestry, so we examine how models trained on ADNI, generalize to a new population dataset from India (the NIMHANS cohort). We first benchmark our models by predicting “brain age” - the task of predicting a person’s chronological age from their MRI scan and proceed to AD classification. We also evaluate the benefit of using a 3D CycleGAN approach to harmonize the imaging datasets before training the CNN models. Our experiments show that classification performance improves after harmonization in most cases, as well as better performance for dMRI as input.

## I. Introduction

Alzheimer’s disease (AD) is the most prevalent age-related neurodegenerative condition, responsible for around 70% of dementia cases globally [1]. At present, about one in three elderly individuals in the United States eventually develop AD or a related type of dementia. As such, there is a pressing need to uncover factors that promote or resist dementia. To assist with this task, approaches are required to objectively diagnose AD at earlier stages, as well as assess disease progression and prognosis. Machine learning and deep learning models offer great potential in AD research, and MRI-based diagnostic tools would also benefit clinical practice, as accurate diagnosis is key to assigning optimal treatments for each patient. In clinical research, automatic classifiers of disease could be used to rapidly screen imaging databases and biobanks to identify genetic or environmental factors that may influence disease progression or resilience. Deep learning can also be directly applied to raw or minimally pre-processed images, bypassing the time-consuming processes of parcellation and quality control involved in traditional region-of-interest methods for brain morphometry. A recent study [2] trained a deep convolutional neural network (CNN) based on the Inception-ResNet-V2 architecture on 85,721 brain MRI scans from 50,876 participants. Through transfer learning and subsequent fine-tuning for AD classification, the model achieved an impressive accuracy of 91.3% in leave-sites-out cross-validation.

In 2020, Wen et al. reviewed [3] over 30 papers that applied convolutional neural networks to classify patients with AD relative to matched healthy controls, using neuroimaging data. They noted two limitations in the field(1)data leakage leading to inflated accuracy estimates, and (2) insufficient testing of models on diverse cohorts and data from different scanners. With a few recent exceptions [4,5], most CNNs developed for detecting rely on T1-weighted brain MRI, the most commonly collected type of brain scan. Nevertheless, others [6,7] have found initial evidence that dMRI, which is more sensitive than T1-weighted MRI to subtle alterations in the brain’s white matter microstructure, can provide metrics strongly correlated with age, dementia severity, and even levels of brain amyloid—a primary contributor to AD pathology not detectable on standard MRI. Here, we set out to investigate the potential advantages of dMRI for two common AD-related deep learning tasks: brain age estimation and AD classification. Intuitively, training a CNN on diffusion MRI (dMRI) might yield superior results than training solely on T1-weighted MRI.

Brain MRI protocols vary considerably, and models trained on data from a specific scanner may perform more poorly when tested on data acquired from a different site or using a different scanning protocol. This ‘domain shift’ problem has prompted the development of domain adaptation methods. These techniques aim to mitigate the differences between multisite brain MRI datasets by adjusting some properties of the testing data to match a reference dataset or the training data. By doing so, domain adaptation improves the generalizability of machine learning models when tested across diverse imaging protocols and acquisition sites. Various Generative Adversarial Network (GAN)-based approaches have been used for medical image harmonization. Liu et al. [8] used style transfer techniques to adjust brain MRI scans to match a designated reference dataset. Sinha et al. [9] found that AD classification improved after using attention-guided GANs for MRI harmonization. Dinsdale et al. [10] included an adversarial subnetwork to predict scan sites from the predictors used for classification, and defeating this adversary generated site-invariant features for classification. Zhao et al. [11] showed that adversarial subnetworks could also adjust for confounding effects of age and sex as well as site, as these features can also be confounded with site and lead to classifiers that do not generalize well. Here, we also examined GAN-based strategies to address domain differences and make our predictive models more robust. We used a 3D unsupervised CycleGAN model developed by Komandur et al. [12] to harmonize the two datasets. We compared the results for both tasks on harmonized and non-harmonized data acquired at the NIMHANS center (India).

## II. Data AND Preprocessing

Diffusion tensor imaging (DTI) – the most widely-used model of brain tissue microstructure – approximates the local diffusion process through a spatially-varying diffusion tensor (principally a 3D Gaussian approximation). Although the tensor model slightly simplifies the diffusion model and higher-order models (NODDI [13], MAP-MRI [14]) are used in research, diffusion tensors can be computed from single-shell dMRI, which is faster and most convenient to collect clinically. Four standard measures are typically used to characterize the diffusion process: fractional anisotropy (FA), and mean, axial, and radial diffusivity (MD, AxD, and RD), which quantify the shape of the tensors at each voxel. These measures are computed from the three principal eigenvalues of the tensor (**Figs. 1** and equations 1 to 4), indicating the primary directions of water displacement or diffusion at each voxel. FA summarizes the directionality of diffusion, while MD is sensitive to cellularity, edema, and necrosis. Radial diffusivity is increased by processes such as demyelination or dysmyelination. Changes in axonal diameters or density also impact radial diffusivity. AxD is also increased by white matter pathology. We evaluate each of these DTI metrics in comparison to T1w images.

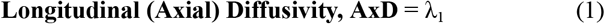

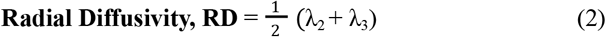

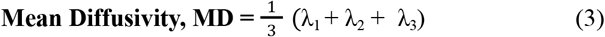

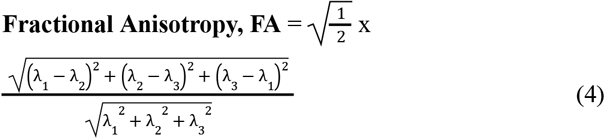

The standard formulae for calculating the diffusion metrics using the diffusion tensor eigenvalues are given in (1) to (4). To evaluate the performance of our deep learning techniques, we first trained and tested CNNs on the task of predicting brain age. In this work, we focus on CNNs as they are widely used and well understood, but other networks (such as vision transformers [15]) have also been used for these and related tasks. While a person’s chronological age is typically known and may not hold immediate clinical utility, predicting brain age in healthy control subjects is a standard benchmarking task, as ground truth is known. When such a trained model is applied to patients at varying stages of dementia, the disparity between the predicted age (referred to as the individual’s “BrainAge”) and their actual chronological age has been associated with subsequent clinical deterioration, the progression of dementia, and even mortality [16].

**Fig. 1.**
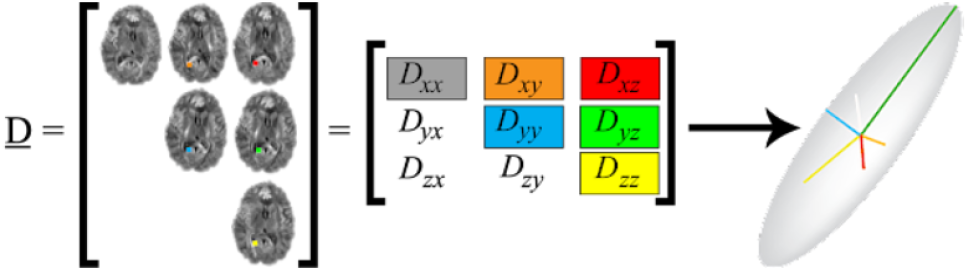
Diffusion tensor components. Tensor components are fitted to the raw diffusion-weighted MRI data, and the eigenstructure of the tensor (right) is used to compute summary metrics of diffusion based on the eigenvalues (the lengths of the 3D vectors in the ellipsoid)

The primary dataset for our experiments is the widely-used, publicly available Alzheimer’s Disease Neuroimaging Initiative (ADNI) dataset – a multisite study launched in 2004 designed to improve clinical trials for the prevention and treatment of Alzheimer’s disease [17]. We used data from a total of 1,195 participants (age: 74.36 ±7.74 years; 600 F/595 M), who had both structural T1w as well as dMRI with a distribution of (633 CN/421 MCI/141 dementia) for our analysis. The second dataset comes from an Indian population assessed at NIMHANS in Bangalore, India [18,19] – a population not typically well represented in neuroimaging studies. This cohort had 301 participants (age: 67.23 ±7.86 years; 169 F/132 M) with a distribution of (123 CN/88 MCI/90 AD). We also analyzed MRIs from 10,000 CN subjects from the UK Biobank [20] for training for the Brain Age experiment, and this had a distribution of age: 64.11 ±7.42 years (5050 F/4950 M).

T1w brain MRI volumes were pre-processed using the following steps [21]: nonparametric intensity normalization (N4 bias field correction), ‘skull stripping’ for brain extraction, nonlinear registration to a template with 6 degrees of freedom and isometric voxel resampling to 2 mm. The pre-processed images were of size 91x109x91. The T1w images were scaled using min-max scaling to take values between 0 and 1. All T1w images were aligned to a common template provided by the ENIGMA consortium [22]. dMRI were non-linearly registered to the T1w images. The dMRI processing pipeline is extensively detailed in [6,7,23].

## III. Deep Learning Architectures

The 3D CycleGAN model [12] (**Fig. 2**) used for harmonization consists of two generators (*G*_*X*_: *X*→*Y* and *G*_*Y*_: *Y*→*X*) and two discriminators (*D*_*X*_ and *D*_*Y*_) corresponding to the source domain *X* (ADNI) and target domain *Y* (NIMHANS). The primary objective was to generate an image representative of the target distribution, when provided an image from the source domain distribution. To achieve this, the model employed an adversarial GAN loss in conjunction with a cyclic consistency loss term. Additionally, it incorporated a patch-based discriminator and identity loss to regularize model training. During training, the model was applied to both pre-processed T1w and DTI map scans obtained from both the source and target datasets separately. The Adam optimizer [24] was used to train both generators and discriminators, with a learning rate of 1x10^-4^ and a batch size of 4. We trained the model for 100 epochs using a multi-step learning rate scheduler with a gamma of 0.1, and steps occurring at 35 and 75 epochs. Overall, the model had 16 million (M) parameters. Each generator and discriminator comprised 5M and 3M parameters, respectively.

**Fig. 2.**
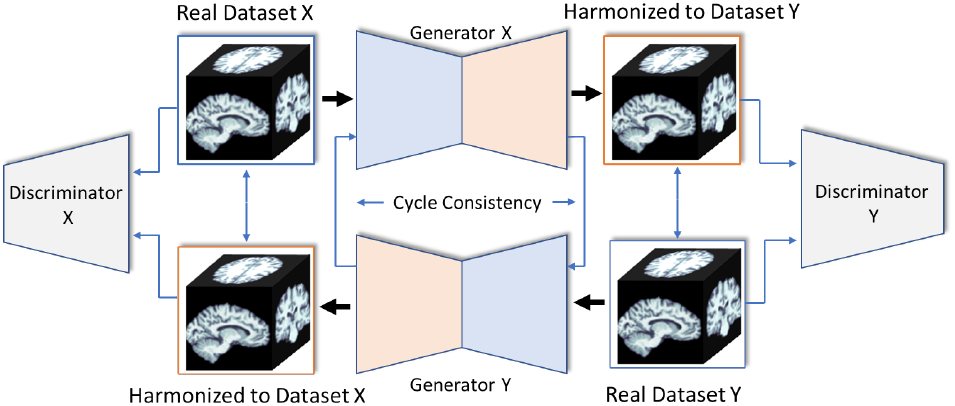
The CycleGAN architecture, with two generators and two discriminators, reproduced from [12].

The 3D CNN architecture (**Fig. 3**) consisted of four 3D Convolution layers with a 3x3 filter size, followed by one 3D Convolution layer with a 1x1 filter, and a final Dense layer. All layers used the ReLu activation function and Instance Normalization. Dropout layers, with a dropout rate of 0.5, and a 3D Average Pooling layer with a 2x2 filter size were added to the 2^nd^, 3^rd^, and 4^th^ layers. Models were trained with a learning rate of 10^-4^, and test performance was assessed using balanced accuracy. We trained this CNN model for 100 epochs, with a batch size of 8. The learning rate was exponentially decayed with a decay rate of 0.96. The Adam optimizer [24] and mean square error loss function were used for training. To deal with overfitting, dropout between layers and early stopping were used. Test performance on Brain age was assessed using the mean absolute error (MAE) to compare results for different modalities. After registering the images to a common template, the data was split into independent training, validation, and testing sets in the ratio of approximately 70:20:10. The 3D-CNN model was first pretrained on T1w scans from 10k subjects in the UK Biobank cohort for age prediction. This pretrained model was then used to predict age in the ADNI and NIMHANS cohort and model performance was assessed. For the AD/CN classification task, the final Dense layer of the model had a sigmoid activation function and the hyperparameters were tuned using grid search.

**Fig. 3.**
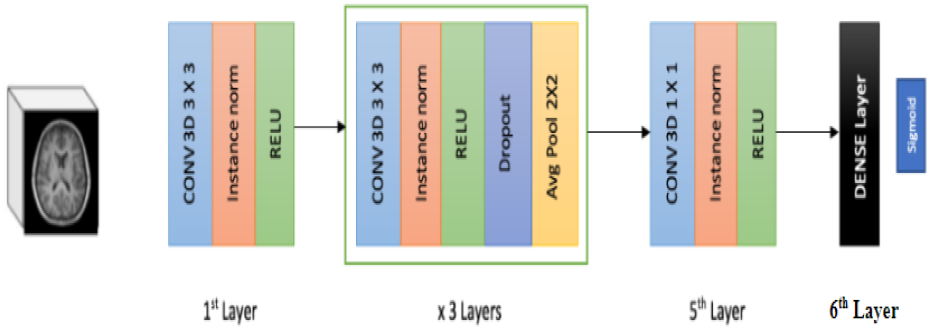
3D CNN Architecture used in the work..

For the dual modality experiments, where two different modality inputs were fed in simultaneously, a concatenated model architecture was used. The 3D CNN architecture (**Fig. 4**) consisted of four 3D Convolutional layers. After flattening, the outputs of these layers were concatenated and passed through a Dense Layer with a sigmoid activation function. This Y-shaped architecture uses distinct CNNs to extract predictive features separately from the anatomical MRI and dMRI. Subsequently, these features are merged for disease classification purposes. The 3D-CNN was trained for 100 epochs with the Adam optimizer, incorporating weight decay regularization. A batch size of 4 was used, and the training process was halted when the validation loss failed to demonstrate improvement for 10 consecutive epochs.

**Fig. 4.**
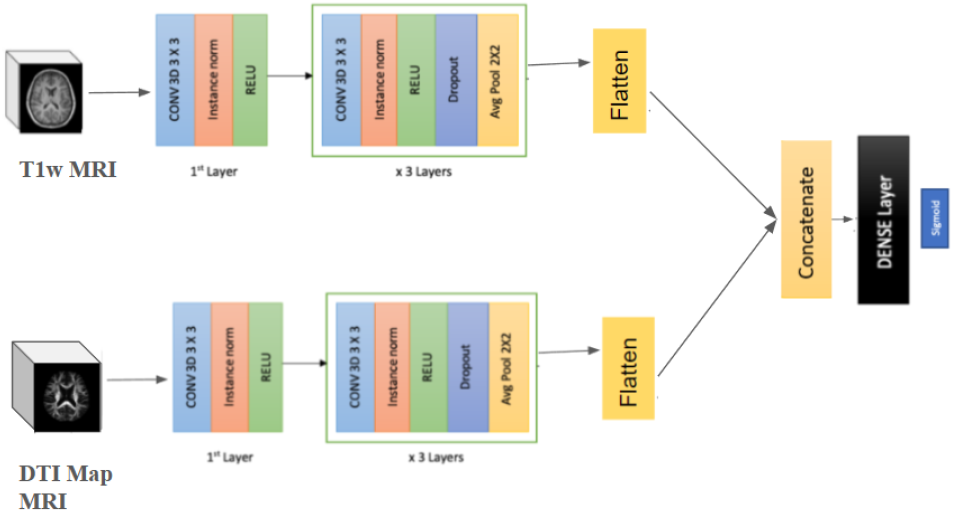
The Concatenated (‘Y-shaped’) 3D CNN Architecture.

## IV. Experimental Results

In the first experiment, we trained the model on the UK Biobank cohort CN subjects’ T1w scans in increasing sample sizes of 2,000 to 10,000, and then used these trained models to predict individual age for ADNI, NIMHANS and harmonized NIMHANS CN subjects’ MRIs separately. The MAE results are shown in **Fig. 5** (note that despite this quite poor performance, subsequent experiments improve on this baseline).

**Fig. 5.**
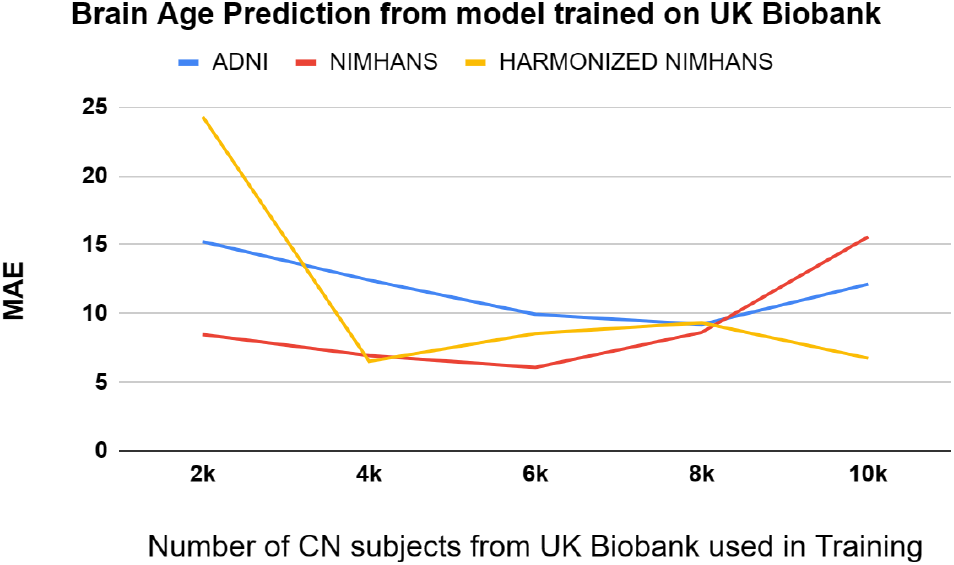
Graph of MAEs for results of training on increasing buckets of UK Biobank CN subjects. The column heads represent the number of subjects in the bucket.

The results show that the distribution of NIMHANS is closer to UK Biobank than ADNI, but both these models need to be finetuned on the respective datasets to improve performance. There may be evidence for an overfitting point for the trained model, which is around 6k subjects, after which the MAEs on the test datasets start increasing again. Harmonization brings the distribution of NIMHANS dataset closer to ADNI.

In the second experiment, we trained the model on ADNI CN subjects, and tested the generalizability of the model on NIMHANS, as a separate test cohort. We trained the model for 100 epochs; MAE results are shown in **Table I**.

**TABLE I.** Results of brain age prediction for a 3D CNN model trained on ADNI CN data. For both datasets, the diffusion MRI methods perform best, although all metrics perform poorly for the NIMHANS dataset.

|  | T1 | DWI-FA | DWI-MD | DWI-RD | DWI-AxD |
| --- | --- | --- | --- | --- | --- |
| ADNI | 5.63 | **3.39** | 4.72 | 4.37 | 4.13 |
| NIMHANS | 9.77 | 8.16 | 7.91 | 18.80 | **7.80** |

The results show that model performance is generally better when DWI-derived maps are used as inputs as compared to the T1w images. The MAEs for models trained on the ADNI dataset were notably lower compared to the corresponding dMRI modalities from the NIMHANS dataset. This performance gap shows the need to train the model on a more diverse range of data to enhance the model’s ability to generalize across different datasets and settings, as in our next experiment.

In the third experiment, we trained the model on a combination of ADNI and NIMHANS CN subjects. The model was then tested separately on a hold out ADNI and NIMHANS test set. MAE results for the experiment are as shown in **Table II**. The experiments were repeated for the ADNI and the harmonized NIMHANS dataset, as shown in **Table II**.

**TABLE II.** Results of brain age prediction for a 3D CNN model trained on combined ADNI and NIMHANS CN data.

|  | T1 | DWI-FA | DWI-MD | DWI-RD | DWI-AxD |
| --- | --- | --- | --- | --- | --- |
| **Trained on ADNI and NIMHANS** |  |  |  |  |  |
| ADNI | 6.28 | **3.87** | 5.77 | 6.13 | 5.11 |
| NIMHANS | 6.56 | **4.27** | 6.27 | 6.59 | 6.14 |
| **Trained on ADNI and harmonized NIMHANS** |  |  |  |  |  |
| ADNI | 6.43 | **3.43** | 4.92 | 4.61 | 4.89 |
| Harmonized NIMHANS | 4.48 | 4.95 | 6.76 | **4.16** | 4.37 |

We compared the performance of our model with two different models from the literature - Peng et al. [25] and Yin et al. [26]. The Peng et al. model was originally trained on data from UK Biobank T1ws, whereas the Yin et al. model was originally trained on ADNI T1ws. On the same three test sets of T1ws as our experiments, using trained model weights without any re-training, the performance is shown in **Table III**.

**TABLE III.**
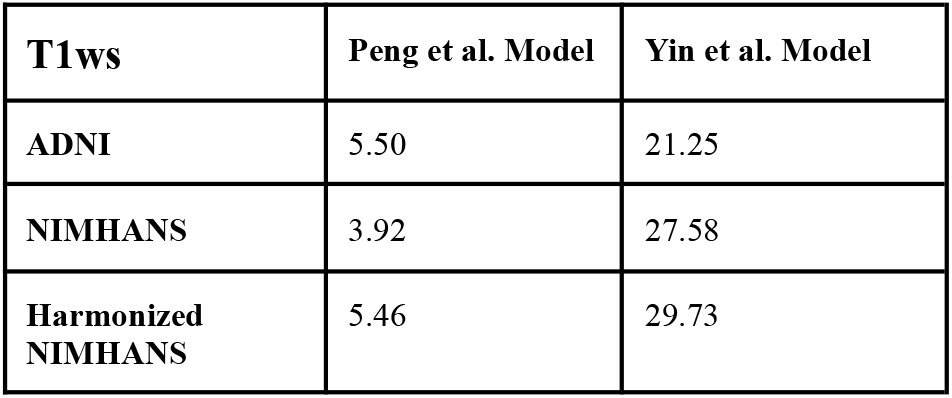
Results of brain age prediction from models from literature.

The best task performance was achieved when DWI-FA was used as the input. This was found for both test datasets in the case of training on ADNI and NIMHANS, with an MAE generally lower than when using other input maps. The performance of the model is better on the test NIMHANS dataset, as compared to results from the second experiment as would be expected with the inclusion of data from the cohort in training. So, despite the model generalizing better, the overall performance of the model is close to the results for the best brain age prediction models as seen in **Table III**. One caveat is that those models are trained on larger datasets, so as more data is provided for training, the performance may improve. Performance improves when the harmonized NIMHANS dataset is used during training with ADNI. In most cases, the MAE goes down. The lowest MAE for ADNI is obtained when the DWI-FA maps are used as inputs, in this case.

In the fourth experiment, we used the concatenated model (**Fig. 4**) to predict Brain Age. We trained the model separately for combinations of T1 and DWI image inputs, for both harmonized and non harmonized NIMHANs along with ADNI separately. The MAE results for the experiments are shown in **Table IV**.

**TABLE IV.** Results of brain age prediction for the Concatenated 3D CNN Model.

|  | T1 +<br>DWI-FA | T1 +<br>DWI-MD | T1 +<br>DWI-RD | T1 +<br>DWI-AxD |
| --- | --- | --- | --- | --- |
| Trained on ADNI and NIMHANS |  |  |  |  |
| ADNI | 11.97 | **4.94** | 5.11 | 5.18 |
| NIMHANS | 12.64 | **4.75** | 5.01 | 5.11 |
| Trained on ADNI and harmonized NIMHANS |  |  |  |  |
| ADNI | 6.31 | 5.13 | **4.73** | 4.85 |
| Harmonized NIMHANS | 6.32 | 5.90 | 6.10 | **4.95** |

Performance on this task is better for the concatenated model, than when using T1w alone, except for the case of combining T1w and DWI-FA. Model performance also improves in most cases when the harmonized NIMHANS data is used, especially for ADNI. This performance is comparable to that of DWIs used alone as input for the models. These results may further improve as larger and more diverse datasets are used for training.

In the fifth experiment, we trained the model on a combination of ADNI and NIMHANS subjects to classify unseen individuals as having CN vs Alzheimer’s disease. We used the balanced accuracy and F1 score metrics to compare the performance for the different input data types (**Table V**). Experiments were repeated for the ADNI and harmonized NIMHANS dataset (**Table V)**.

**TABLE V.** Results of CN/AD classification for a 3D CNN model trained on combined cohort data.

|  | T1 | DWI-FA | DWI-MD | DWI-RD | DWI-AxD |
| --- | --- | --- | --- | --- | --- |
| Trained on ADNI and NIMHANS |  |  |  |  |  |
| ADNI |  |  |  |  |  |
| Balanced Accuracy | 0.62 | 0.83 | **0.87** | 0.86 | **0.87** |
| F1 Score | 0.64 | 0.80 | **0.84** | 0.83 | **0.84** |
| NIMHANS |  |  |  |  |  |
| Balanced | 0.63 | **0.90** | 0.84 | 0.87 | 0.80 |
| Accuracy |  |  |  |  |  |
| F1 Score | 0.59 | **0.87** | 0.78 | 0.82 | 0.74 |
| Trained on ADNI and harmonized NIMHANS |  |  |  |  |  |
| ADNI |  |  |  |  |  |
| Balanced Accuracy | 0.49 | 0.75 | 0.74 | **0.74** | 0.72 |
| F1 Score | 0.89 | 0.68 | 0.68 | **0.71** | 0.64 |
| Harmonized NIMHANS |  |  |  |  |  |
| Balanced Accuracy | 0.76 | 0.91 | 0.75 | **0.93** | 0.91 |
| F1 Score | 0.95 | 0.94 | 0.98 | **0.93** | 0.90 |

For non-harmonized NIMHANS and ADNI, the performance is better for the DWI-derived maps used as inputs, as compared to the T1w images. The best Balanced Accuracy and F1 Score is for DWI-MD for ADNI and DWI-FA for NIMHANS respectively. Harmonizing the NIMHANS dataset increases the balanced accuracy for most types of input data in the AD classification task. The performance was also slightly better for the trained model on the holdout NIMHANS dataset, compared to the ADNI holdout test dataset. The results for DWI maps are comparable to those obtained from T1w using larger sample sizes from ADNI, as reported in [27]. The model performance for T1w inputs might improve with more training datasets, but the results suggest that a smaller dataset of DWI modalities is a better substitute for the AD classification task.

In the sixth experiment, we used the concatenated model (**Fig. 4**) to classify people with AD versus CN. We trained the model separately for combinations of T1 and DWI-derived maps as inputs, for both harmonized and non harmonized NIMHANS, along with ADNI separately. The results of the experiment are shown in **Table VI**.

**TABLE VI.**
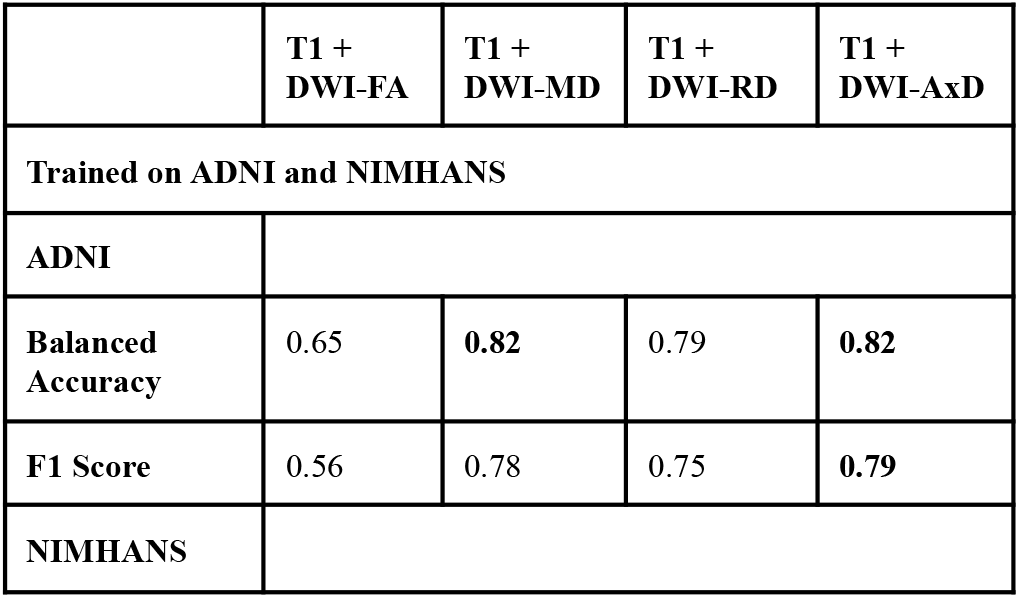
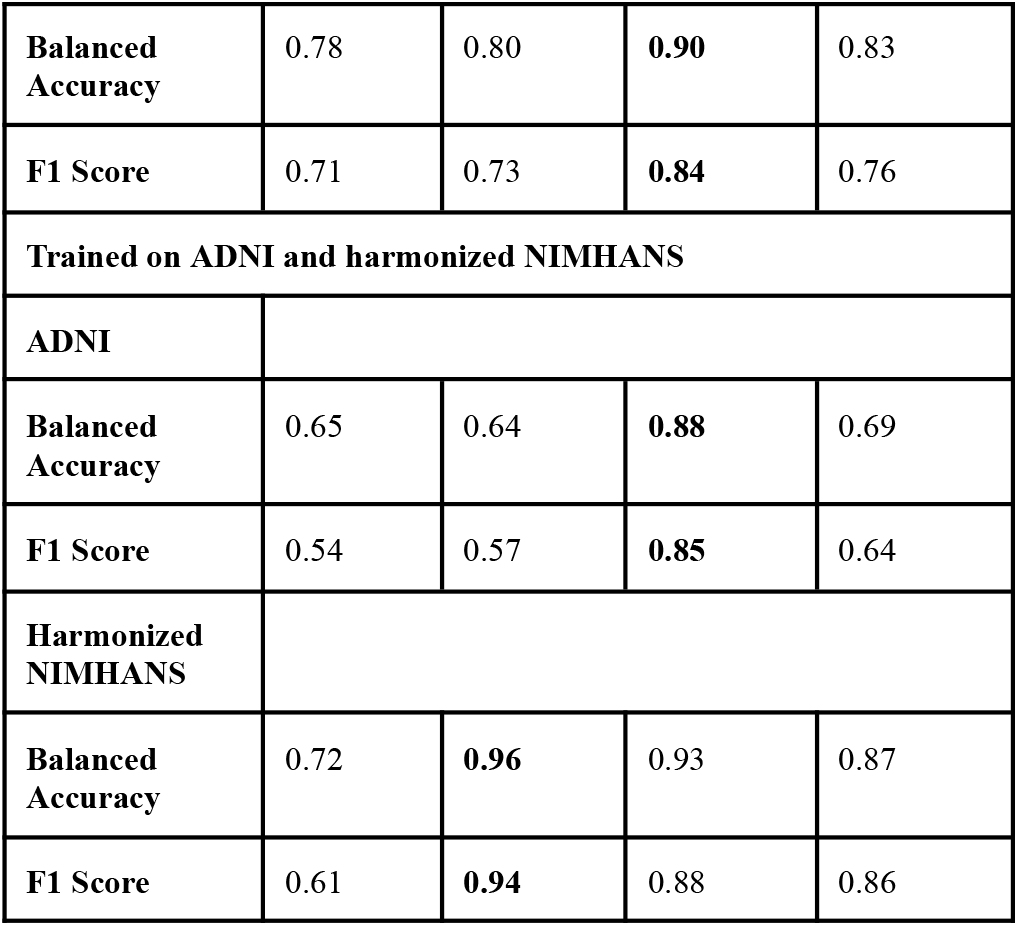
Results of CN/AD classification for the concatenated CNN model.

Task performance was better for the concatenated model, than when using T1w input data alone. Model performance also improves in most cases when the NIMHANS data is harmonized. In most cases, the performance is comparable to that of DWIs used alone as input for the models. Larger or more diverse training data may further improve model performance. The best performance was observed when T1w and DWI-AxD maps were combined. The worst performance was in the case of T1w and DWI-FA images concatenated as inputs. The performance was also slightly better for the trained model on the holdout NIMHANS dataset, as compared to the ADNI holdout test dataset. Again, these results are comparable to those obtained when training on larger samples of the ADNI dataset [27].

We also visualized saliency maps using Grad-CAM [28] for the best performing model in both experiments for Brain Age prediction and CN vs Alzheimer’s Disease Classification. In **Fig. 6**, the first column shows the input image, the second column shows the regions contributing to model performance and the last column shows these two images superimposed. We show saliency maps for the best performing modality from the two experimental setups.

**Fig. 6.**
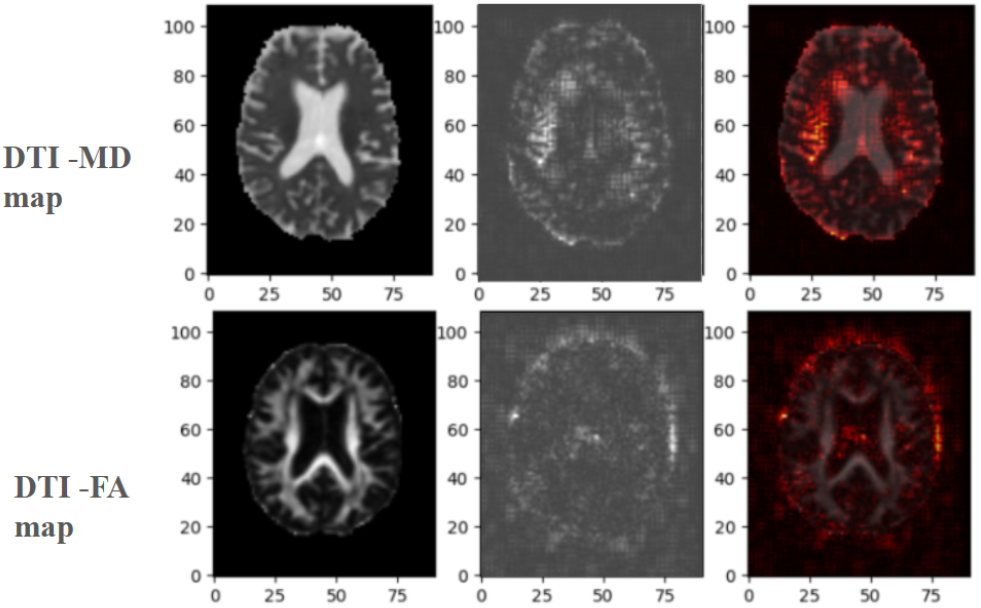
Saliency maps for both tasks for the best performing input types.

## V. Conclusion AND Future Work

Using diverse datasets, we benchmarked CNN models for two common tasks in dementia research - brain age prediction and AD classification. We examined how well a model trained on an open dataset generalized to a cohort of different ancestry that was not seen during training; we also assessed performance improvements when the cohorts are mixed to create the training datasets. In general, performance improved after harmonizing the datasets using a deep learning model based on unsupervised 3D CycleGAN. Training the models on a combined dataset featuring diverse cohorts also improved performance on these tasks. When smaller datasets are available for training, AD classification was more accurate when based on DWI-derived maps, compared to T1w images. DWI image modalities may complement T1w images for other prediction tasks [28,29]. A concatenated model with multiple image modality inputs also outperformed a model using only T1w MRIs.

In future, we plan to include matched T1w and DWI data, from larger, more diverse datasets, scanned with additional protocols, to comprehensively evaluate model performance. Performance may improve with more advanced diffusion MRI metrics, such as TDF-FA (which computes a tensor distribution function from single-shell diffusion MRI, and weights the FA appropriate when multiple crossing fibers are detected within a voxel). We will also evaluate other image harmonization techniques including StyleGANs [9], VAE-GANs [30], or CALAMITI [31], to improve out-of-domain generalizability. These approaches can make data from diverse scanners and protocols more comparable, mitigating domain shift.

## ACKNOWLEDGMENTS

This work was supported by NIH grant R01 AG060610. NIMHANS data collection was supported by the Department of Science and Technology, Govt. of India, grant nos. DST-SR/CSI/73/2011 (G); DST-SR/CSI/70/2011 (G); and DST/CSRI/2018/249 (G).

## References

[1] World Health Organization, “Dementia,” 2022. https://www.who.int/news-room/fact-sheets/detail/dementia

[2] Lu, B., Li, H. X., et.al. (2022). A practical Alzheimer’s disease classifier via brain imaging-based deep learning on 85,721 samples. Journal of Big Data, 9(1), Article 101.

[3] Wang L., Pei J., Zhan Y., Wen Y., “Overview of Meta-Analyses of Five Non-pharmacological Interventions for Alzheimer’s Disease,” Front. Aging Neurosci. 12:594432 (2020).

[4] Wang D., et al., “Application of multimodal MR imaging on studying Alzheimer’s disease: a survey,” Curr. Alzheimer Res. 877–92 (2013).

[5] Knudsen L., et.al., “The role of multimodal MRI in mild cognitive impairment and Alzheimer’s disease,” J Neuroimaging 148–157 (2022).

[6] Thomopoulos S., et al., “Diffusion MRI Metrics and their relation to Dementia Severity: Effect of Harmonization Approaches,” (2021) In 17th International Symposium on Medical Information Processing and Analysis (Vol. 12088, pp. 166–179). SPIE.

[7] Chattopadhyay, T., et al. (2023). Predicting dementia severity by merging anatomical and diffusion MRI with deep 3D convolutional neural networks. In the 18th International Symposium on Medical Information Processing and Analysis (SIPAIM; Vol. 12567, pp. 90–99). SPIE.

[8] Liu, M., et.al. (2021). Style transfer using generative adversarial networks for multi-site MRI harmonization. MICCAI 2021 (pp. 313–322).

[9] Sinha, S., et.al. (2021). Alzheimer’s disease classification accuracy is improved by MRI harmonization based on attention-guided generative adversarial networks. In the 17th International Symposium on Medical Information Processing and Analysis (Vol. 12088, pp. 180–189).

[10] Dinsdale, N. K., Jenkinson, M., & Namburete, A. I. (2021). Deep learning-based unlearning of dataset bias for MRI harmonisation and confound removal. NeuroImage, 228, 117689.

[11] Zhao, Qingyu et al. “Training confounder-free deep learning models for medical applications.” Nature communications vol. 11,1 6010. 26 Nov. 2020.

[12] Komandur, D., et al. (2023). Unsupervised harmonization of brain MRI using 3D CycleGANs and its effect on brain age prediction. 19th International Symposium on Medical Information Processing and Analysis (SIPAIM) (pp. 1–5). IEEE.

[13] Deligianni, F., et al. (2016). NODDI and tensor-based microstructural indices as predictors of functional connectivity. Plos one, 11(4), e0153404.

[14] Özarslan, E., Koay, C. G., et.al. (2013). Mean apparent propagator (MAP) MRI: a novel diffusion imaging method for mapping tissue microstructure. NeuroImage, 78, 16–32.

[15] Dhinagar, N. J., Thomopoulos, S. I., Laltoo, E., & Thompson, P. M. Efficiently Training Vision Transformers on Structural MRI Scans for Alzheimer’s Disease Detection. ArXiv.

[16] Cole, J. H., et al. (2018). Brain age predicts mortality. Molecular Psychiatry, 23(5), 1385–1392.

[17] Veitch D., et al., “Understanding disease progression and improving Alzheimer’s disease clinical trials: Recent highlights from the Alzheimer’s Disease Neuroimaging Initiative,” Alzheimer’s Dement. 15(1):106–152 (2019).

[18] Bharath, S., et.al. (2017). A Multimodal Structural and Functional Neuroimaging Study of Amnestic Mild Cognitive Impairment. American Journal of Geriatric Psychiatry, 25(2), 158–169.

[19] Joshi, H., et.al. (2019). Differentiation of Early Alzheimer’s Disease, Mild Cognitive Impairment, and Cognitively Healthy Elderly Samples Using Multimodal Neuroimaging Indices. Brain Connectivity, 9(9), 730–741.

[20] Sudlow, C., et al. (2015). UK Biobank: An open access resource for identifying the causes of a wide range of complex diseases of middle and old age. PLoS Medicine, 12(3), e1001779.

[21] Lam, P., et al., “3-D Grid-Attention Networks for Interpretable Age and Alzheimer’s Disease Prediction from Structural MRI,” (2020).

[22] Jahanshad, N., et. al. (2013). Multi-site genetic analysis of diffusion images and voxelwise heritability analysis: a pilot project of the ENIGMA-DTI working group. NeuroImage, 81, 455–469.

[23] Feng, Y., et.al. “Deep Normative Tractometry for identifying joint white matter Macro- and Microstructural abnormalities in Alzheimer’s Disease.” EMBC 2024.

[24] Kingma D., Ba J. (2015). “Adam: A method for Stochastic Optimization,” ICLR (2015).

[25] Peng, H., Gong, W., Beckmann, C. F., Vedaldi, A., & Smith, S. M. (2021). Accurate brain age prediction with lightweight deep neural networks. Medical image analysis, 68, 101871.

[26] Yin, Chenzhong, et al. ‘Anatomically interpretable deep learning of brain age captures domain-specific cognitive impairment.’ Proceedings of the National Academy of Sciences 120.2 (2023):e2214634120.

[27] Dhinagar, N. J., et al. (2021). 3D convolutional neural networks for classification of Alzheimer’s and Parkinson’s disease with T1-weighted brain MRI. In the 17th International Symposium on Medical Information Processing and Analysis (Vol. 12088, pp. 277–286). SPIE.

[28] H. Zhao, et al. Deep learning based diagnosis of Parkinson’s Disease using diffusion magnetic resonance imaging.” Brain Imaging and Behavior vol. 16, 4 (2022): 1749–1760.

[29] Chattopadhyay, T., et. al. (2023). Comparison of Anatomical and Diffusion MRI for detecting Parkinson′ s Disease using Deep Convolutional Neural Network. EMBC 2023.

[30] Moyer D., Ver Steeg G., Tax C., Thompson P.M., “Scanner invariant representations for diffusion MRI harmonization,” Magn Res Med. 84(4):2174–2189 (2020).

[31] Zuo, L., et. al. (2021). Unsupervised MR harmonization by learning disentangled representations using information bottleneck theory. NeuroImage, 243, 118569.

[32] Selvaraju, R. R., et al. (2017). Grad-cam: Visual explanations from deep networks via gradient-based localization. In Proceedings of the IEEE international conference on computer vision (pp. 618–626).

[33] Lam, P. K., et al., “Accurate brain age prediction using recurrent slice-based networks,” In 16th International Symposium on Medical Information Processing and Analysis (Vol. 11583, pp. 11–20). SPIE.(2020).

[34] Gupta Y., Kim J., Kim B., Kwon Goo-Rak, “Classification and Graphical Analysis of Alzheimer’s Disease and Its Prodromal Stage Using Multimodal Features From Structural, Diffusion, and Functional Neuroimaging Data and the APOE Genotype”, Front Aging Neuroscience 12:238 (2020).

[35] Gupta, U., Chattopadhyay, T., Dhinagar, N., Thompson, P. M., & ver Steeg, G. (2023). Transferring Models Trained on Natural Images to 3D MRI via Position Encoded Slice Models. In 2023 IEEE 20th International Symposium on Biomedical Imaging (ISBI) (pp. 1–5). IEEE.

[36] Gupta, U., Lam, P., ver Steeg, G., et al., “Improved Brain Age Estimation with Slice-based Set Networks,” In 2021 IEEE 18th International Symposium on Biomedical Imaging (ISBI) (pp. 840–844).

[37] Nir T., Jahanshad N., Villalon, J., Isaev D., Thompson P.M., “Fractional anisotropy derived from the diffusion tensor distribution function boosts power to detect Alzheimer’s disease deficits,” Magn Reson Med. 78(6):2322–2333 (2017).

[38] Lin, W., Tong, T., Gao, Q., et al., “Convolutional Neural Networks-Based MRI Image Analysis for the Alzheimer’s Disease Prediction From Mild Cognitive Impairment,” Front. Neurosci. 12:777 (2018).

[39] Kochunov P., Hong L., Dennis E., Morey R., Tate D., Nir T., Glahn D., Thompson P.M., Jahanshad N., “ENIGMA-DTI: Translating reproducible white matter deficits into personalized vulnerability metrics in cross-diagnostic psychiatric research,” Human Brain Mapping 194–206 (2022).

[40] Mirzaalian H., Ning L., Savadjiev P., et al., “Inter-site and inter-scanner diffusion MRI data harmonization,” NeuroImage 135:311–23 (2016).

